# Synaptic degeneration in neuronal circuits hinders memory recall, memory rescue, and learning

**DOI:** 10.1101/2022.08.22.504766

**Authors:** Kwan Tung Li, Daoyun Ji, Changsong Zhou

**Affiliations:** Department of Physics, Centre for Nonlinear Studies, Beijing–Hong Kong–Singapore Joint Centre for Nonlinear and Complex Systems (Hong Kong), Institute of Computational and Theoretical Studies, Hong Kong Baptist University, Hong Kong; Department of Neuroscience, Baylor College of Medicine, Houston, TX 77030, USA; Department of Molecular and Cellular Biology, Baylor College of Medicine, Houston, TX 77030, USA

## Abstract

Neurodegeneration is a characteristic of Alzheimer’s disease (AD), but its effect on neural activity dynamics underlying memory deficits is unclear. Here, we studied the effects of synaptic degeneration on neural activities associated with memory recall, memory rescue by slow-gamma stimulation, and learning a new memory, in an integrate-and-fire neuronal network. Our results showed that reducing connectivity decreases the neuronal synchronisation of memory neurons at slow-gamma frequencies and impairs memory recall performance. Although slow-gamma stimulation rescued memory recall and slow-gamma oscillations, the rescue with reduced connectivity caused a side effect of activating mixed memories. During the learning of a new memory, reducing connectivity caused an impairment in storing the new memory, but did not affect previously stored memories. Our results reveal potential computational mechanisms underlying the memory deficits caused by synaptic degeneration in AD.

## Introduction

Abnormal amyloid-beta (Aβ) plague deposition is a key feature of Alzheimer’s disease (AD) and they affect brain circuits at different levels (Palop et al., 2007; Palop and Mucke, 2016, 2010; Shankar et al., 2007). At the neuronal level, action potential peaks are reduced (Kerrigan et al., 2014). At the synapse level, synaptic currents are potentiated at low concentrations of Aβ, but depressed at high concentrations (Puzzo et al., 2008). At the circuit level, Aβ induces synaptic degeneration, leading to a reduction in the probability of synaptic connectivity between neurons (Li et al., 2009; Pascale et al., 2007; Shankar et al., 2007). This reduced connectivity may change the dynamic state of a circuit and affect its responses to external stimuli (Denève and Machens, 2016). In some mouse models of AD, excitatory neurons are over-excited and the excitation–inhibition (E-I) balance breaks down, leading to deficits in spatial memory performance (Busche et al., 2015). In other animal models, however, excitatory neurons are under-excited and hippocampal slow-gamma (~40 Hz) oscillations are reduced, which also produces spatial memory deficits (Etter et al., 2019). How these seemingly contradictory results in circuit excitability and their intricate relationship with the degeneration of connectivity at different degrees are not clear.

Memory deficits in AD may be a result of impaired memory recall or impaired learning, leading to improper memory storage. Impaired memory recall is likely to be associated with the activation of memory engrams (Roy et al., 2016; Ryan et al., 2015). Optogenetic activation of engram cells in the dentate gyrus of the hippocampus restores memory recall in an AD model (Roy et al., 2016), pointing to the involvement of improper memory recall in the memory deficits of AD. This is supported by the rescue of memory recall via slow-gamma optogenetic stimulations in another AD model (Etter et al., 2019). In addition, it is well known that the formation of new memories during learning is severely impaired in AD and AD-related animal models (Cheng and Ji, 2013; Germano and Kinsella, 2005; Hodges et al., 1990; Stopford et al., 2007; Takashima, 2012; Weintraub et al., 2012). However, it is unclear how these different aspects of memory deficits are related to different degrees of connectivity deficit.

In this study, we aimed to understand how connectivity deficits lead to changes in neuronal activity patterns and the relevant neural oscillations that underlie the impaired recall of existing memories, the rescue of memory recall by slow-gamma stimulation, and impaired learning. Because it is difficult to experimentally monitor specific engram cells associated with different memories and their dynamic changes at the neuronal and synaptic levels, we conducted our study using a computationally simulated neural network model, where the degree of synaptic connectivity could be manipulated and neural firing patterns, their dynamics, and associated synaptic changes could be precisely monitored.

We found that reducing connectivity during memory recall caused a transition of memory-coding neurons from a synchronous state to an asynchronous state, reduced slow-gamma oscillations, and reduced neural activities associated with memory recall. Second, our simulation showed that rescue using external slow-gamma stimulation at low connectivity caused a side effect of co-activating multiple memories, despite an improvement in recall performance. Third, during the learning of a new memory in the presence of existing memories, our results showed that reducing connectivity induced an impairment in coding the new memory engram, but did not strongly deteriorate existing memories.

## Results

We started by studying the effects of reduced connectivity on neural activities related to memory recall. The model we used was a randomly connected conductance-based E-I neuronal circuit (Yang et al., 2017) consisting of 2,000 excitatory and 400 inhibitory integrate-and-fire neurons (***Figure 1a***). The interneurons were able to receive external stimulation for memory rescue, but the rescue stimulation was turned off here to study memory recall. We considered a case of 10 stored memories in our model. Each single memory was coded in a preset group of 200 non-overlapping excitatory neurons (memory engram) connected via strong connections, while other connections (between memory engrams) were at a weak baseline level. All connections were randomly assigned a connection probability (connectivity, *C*). We modelled the synaptic degeneration seen in AD by systematically reducing *C* (see Methods).

**Figure 1.**
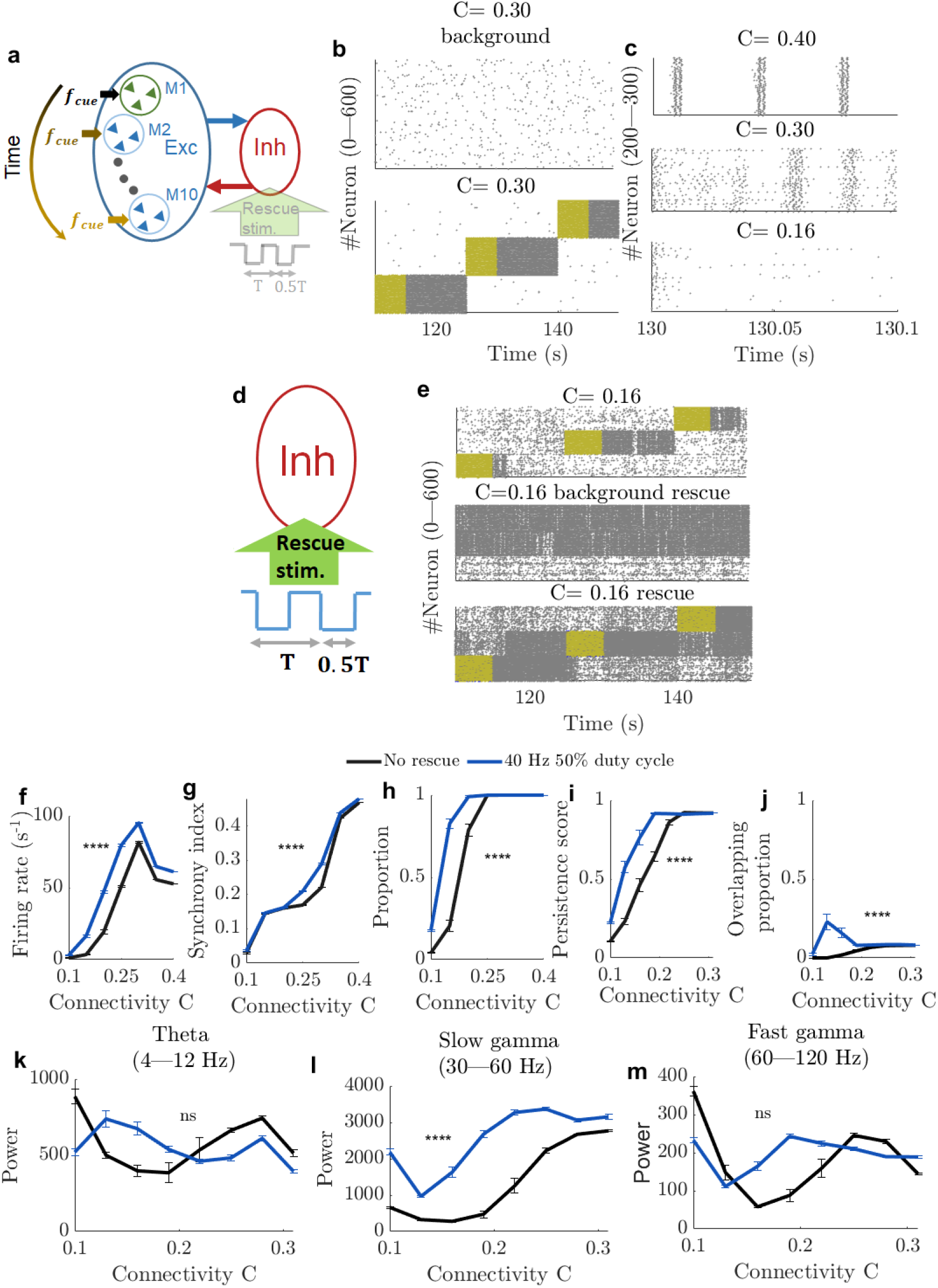
Reduced connectivity impairs memory recall and memory rescue. (a): Schematic of the model. The circuit consisted of excitatory (2,000) and inhibitory (400) neurons. All neurons, both excitatory and inhibitory, were randomly connected with a probability of C and received an external backgrond (2.5 Hz) input to maintain their baseline activity. Within the excitatory population, 10 preset memories (M1–M10) were stored. Memory rescue was not used to study memory recall. (b): Spike raster plots of 400 excitatory neurons (corresponding to two memory engrams) at C = 0.30 in a background state (top) and during memory recall with a strong cue input (bottom). The periods with a strong cue (12.5 Hz) input (5 s) are marked in yellow. Note the intense firing of cued memory neurons (persistent state) even after cue termination. (c): Close-up view of the spike raster plot (100 ms window, 100 neurons in a memory engram) after cue termination at different C values: 0.4 (top), 0.3 (middle), 0.16 (bottom). (d): As in (a), but with memory rescue. Rescue stimulation at 40 Hz (T = 25 ms) with a 50% duty cycle was applied to half of the inhibitory neurons. (e): Spike raster plots of neurons in three memory engrams under different conditions. In this example of memory rescue, C = 0.16 and rescue stimulation (40 Hz, 50% duty cycle) was turned off during the recall with a strong cue input (top), turned on in the background state without the cue (middle), or turned on during recall (bottom). (f–h): Quantifications of cued memory neurons during memory recall with and without rescue stimulation – population mean firing rate (f), synchrony index (g), and proportion of high-firing memory neurons (h). (i–m) Quantifications of persistent states and local field potential oscillations with (red) and without (black) rescue stimulation at different C values – persistence score (i), overlapping proportion (j), theta power (k), slow-gamma power (l), fast-gamma power (m). Error bars: standard error of the mean (SEM). n = 10 trials per p value. ns: p > 0.05; ****: p < 0.0001; two-way analysis of variance (2ANOVA) test.

### Memory Recall

In addition, we modelled memory recall by applying a strong cue input (12.5 Hz Poisson spike trains), *f_cue_*, for 5 s to every neuron in a memory engram (relative to a weak 2.5 Hz background input) and the 10 memory engrams were recalled sequentially. As shown in an example raster plot, the background activity in the absence of the cue input was low (***Figure 1b top***). The cue input often moved (or activated) the neurons in a memory engram into a state characterised by a high level of firing activity that persisted even after the cue was terminated (persistent state), while other neurons remained at a low-activity state (***Figure 1b bottom***). The persistent state is commonly studied in working memory models of spiking neurons (Bi and Zhou, 2020; Brunel and Wang, 2001; Goldman-Rakic, 1995; Mongillo et al., 2008; Wang et al., 2013, 2011; Wang, 1999). Here, we used the cue-triggered persistent states after cue termination as a proxy of neural activity underlying memory recall, because these states are produced by the strong excitatory-to-excitatory connections within a memory engram (Mongillo et al., 2008; Tsodyks, 2005; Wang and Wang, 2001). Therefore, to study how connectivity affects memory recall, we examined how neural activities associated with persistent states depended on *C.*

Based on raster plots at various *C* values (***Figure 1c***), we observed a dynamic change in firing rate after the cue input and firing synchrony among neurons in the cued memory engram (cued memory neurons). When *C* was high (at 0.40, *Figure 1c* top), cued memory neurons fired at a high rate and their spikes were synchronised and phase-locked to oscillations at a frequency of approximately 40 Hz. When *C* was decreased to 0.30 (***Figure 1c*** middle), the firing rates increased, but became less synchronised and less phase-locked. When *C* was further decreased to a low value of 0.16 (***Figure 1c*** bottom), the cued memory neuron firings eventually became non-synchronous and decreased to a very low rate.

We quantified this observation by measuring the firing activities of cued memory neurons in a window from the cue termination to 10 s after termination (before the onset of another cue input to another memory engram). The measurements included the population mean firing rate (number of spikes per neuron within cued memory engram per second), the synchrony index, and the proportion of high-firing neurons (those with a rate > 5 s^−1^, the threshold to separate a low-activity state from a persistent state; see Methods and below) in the cued memory engram. We found that a mild reduction in *C* induced a significant increase in the population mean firing rate. However, once the rate reached a peak, it rapidly decreased with reduction in the *C* value (***Figure 1f*** black line). In contrast to the non-monotonous change in the population mean firing rate, the firing synchrony (***Figure 1g*** black line) and the proportion of high-firing neurons (***Figure 1h*** black line) decreased monotonously as *C* decreased, with steep changes occurring at *C* = 0.25–0.35. Additional analyses found that low *C* values (< 0.2) led to low firing rates and low firing synchrony in a range of synaptic weights connecting excitatory neurons to inhibitory neurons (*g^E→I^* from 0.10 to 0.20), suggesting that the effects of *C* were not sensitive to specific parameter values in our model (***Appendix – figure 1***). These results indicate that neuronal activities associated with memory recall were compromised as *C* was reduced, suggesting that reduced connectivity impairs memory recall.

To further characterise recall performance, we next detected persistent states and quantified how their properties depend on *C*. A persistent state of a cued memory engram was defined as a time period after cue input termination when the mean firing rate of its memory neurons within a 1 s window at each 1 ms time step exceeded a threshold of 5 *s*^−1^ (see Methods). This threshold was determined because the firing rates of all excitatory neurons within 10 s after cue input termination displayed a bimodal distribution when *C* was high (> 0.19), with two groups of neurons separated by a rate of ~5 *s*^-1^ (***Appendix – figure 2A***), whereas those with background activity levels did not, with a single group of low-rate neurons (***Appendix – figure 2B***). We quantified each persistent state using two metrics. The first metric was a persistence score for each persistent state, which was used to quantify how its duration deviated from the expected length (from the termination of the cue input to one memory engram to the beginning of the cue input to another). The persistence score was between 0 and 1, with a larger value indicating a duration closer to the expected length. The second metric was the overlapping proportion, which was used to quantify the degree of co-activation of persistent states of more than one memory engram, when one of the engrams was cued (see Methods). We found that at high *C* values (≥ 0.25) the persistence score was close to 1, while at low *C* values (≤ 0.25) the score decreased rapidly (***Figure 1i***). In contrast, the overlapping proportion was always low (***Figure 1j***). This result suggests that reduced connectivity impairs engram activation.

Given the experimental evidence for the involvement of neural oscillations, especially those with slow-gamma frequencies (30–60 Hz), in memory recall (Etter et al., 2019), we also quantified the power of slow-gamma oscillations associated with memory recall (10 s after cue input termination) in our model, as well as those of theta (4–12 Hz) and high-gamma (60–120 Hz) oscillations for comparison. We found that slow-gamma oscillation power decreased rapidly as *C* was reduced (***Figure 1l***), whereas theta and fast-gamma oscillation power were less sensitive (***Figure 1k, m***). Therefore, our results suggest that the impairment in engram activation at low *C* values is accompanied by an impairment in slow-gamma oscillations.

### Rescue of Memory Recall by Slow-Gamma Stimulation

We next studied how reduced connectivity affects the rescue of memory recall by slow-gamma stimulation. In Ref. (Etter et al., 2019), optogenetic stimulation at 40 Hz with a 50% duty cycle (on 50% and off 50% of the time), intended to suppress hippocampal inhibitory neurons, rescued memory recall performance and restored hippocampal slow-gamma oscillations. To understand the activity dynamics underlying the rescue and the effects of reduced connectivity, we turned on the rescue stimulation in our model (***Figure 1d***). This simulated optogenetic stimulation by resetting the membrane potentials in 50% of inhibitory neurons to their leakage membrane potentials periodically at a frequency of 10–120 Hz. Except for the first 5 s when the circuit was in a transient state starting from random initial conditions, the stimulations were applied throughout the simulation of memory recall, with a duty cycle of up to 90% on (e.g., a 40% duty cycle would mean 40% on and 60% off). We then investigated how rescue stimulation alters the persistent states associated with the recall of stored memories and related neural oscillations at different *C* values.

As in previously reported experiments (Etter et al., 2019), we first applied rescue stimulations at a slow-gamma frequency of 40 Hz with a 50% duty cycle and examined the spike raster plots of neurons in multiple memory engrams. For example, at a low *C* value (0.16), cued memory neurons without rescue stimulation did not always display long-lasting high firing activities after cue input (persistent states), with some terminating early and some missing altogether (***Figure 1e*** top). At the same low *C* value with rescue stimulation, engrams sometimes entered persistent states spontaneously during background activity, in the absence of the cue input (***Figure 1e*** middle). In response to the cue input when rescue stimulations were present, persistent states emerged in most cued memory neurons, although sometimes they lasted longer than expected (***Figure 1e*** bottom). However, the rescue stimulations also produced something peculiar: a persistent state in one memory engram was triggered by the cue input to another engram, or the persistent state lasted too long and was not reset by the cue input to another engram, resulting in the co-activation of multiple memory engrams (***Figure 1e*** bottom), suggesting the mixed recall of multiple memories.

Using the same metrics of memory neurons described above, we compared their activities during memory recall (10 s after cue input termination), with and without rescue stimulation. The population mean firing rates increased significantly after rescue stimulation at all levels of *C* (two-way analysis of variance [2ANOVA], *F*_(1, 158)_ = 270.2, *p* = 1.93 × 10^-33^, ***Figure 1f***). While the synchrony index increased slightly (2ANOVA, *F*_(1, 158)_ = 92.58, *p* = 7.52 × 10^-17^, ***Figure 1g***), the proportion of high-firing neurons was significantly higher after rescue stimulation (2ANOVA, *F*_(1, 158)_ = 46.19,*p* = 3.36 × 10^5^, ***Figure 1h***).

We then compared the metrics of persistent states with and without rescue stimulations. The same threshold of 5 *s*^-1^ was used to detect persistent states when rescue stimulations were applied, because a bimodal firing rate distribution was also present during memory recall (***Appendix – figure 2C***), with the difference that such a bimodal distribution also occurred when rescue stimulations were applied to background activity in the absence of the cue input (***Appendix – figure 2D***). We found that the persistence score was significantly increased by rescue stimulations (2ANOVA, *F*_(1, 78)_ = 75.76, *p* = 7.04 × 10^-15^, ***Figure 1i***). Therefore, our simulation result confirmed the rescue of memory recall by slow-gamma stimulations reported in previous experiments (Etter et al., 2019). In addition, the overlapping proportion, which was low at all *C* values without rescue stimulations, increased, especially for the low-C-value (≤ 0.19) regime (2ANOVA, *F*_(1, 78)_ = 28.83, *p* = 3.26 × 10^-7^, ***Figure 1j***). Thus, our simulation showed that rescue stimulations produce a side effect of co-activating multiple memories during the recall of only one memory. Additional analysis showed that reducing the duty cycle of rescue stimulations to 10%–40% increased the persistence score and decreased the overlapping proportion at low *C* values (***Appendix – figure 3***), suggesting better memory rescue with fewer side effects when slow-gamma stimulations were applied at a lower duty cycle.

We further analysed how rescue stimulations alter neural oscillations. In a previous experiment of memory rescue in mice with AD by 40 Hz with 50% duty duty cycle optogenetic stimulations (Etter et al., 2019), slow-gamma oscillations (30–60 Hz) were enhanced while theta (4–12 Hz) and fast-gamma (60–120 Hz) oscillations were unchanged. Our simulation results were consistent with this experimental finding. When *C* was less than 0.25 and rescue stimulations were applied, there was a significant enhancement in slow-gamma power (2ANOVA, *F*_(1 158)_ = 319.94, *p* = 6.32 × 10^-39^, ***Figure 1l***), but not in theta (2ANOVA, *F*_(1 58)_ = 0.64, *p* = 0.43, ***Figure 1k***) or fast-gamma power (2ANOVA, *F*_(1 58)_ = 2.63, *p* = 0.11, ***Figure 1m***) compared with baseline values.

We sought to determine whether 40 Hz was the optimal stimulation frequency for memory rescue and how the effects of rescue stimulations on persistent states and neural oscillations depended on the stimulation frequency. We found that at a low *C* value (0.16), persistence scores showed the greatest improvement when the frequency was in the range of 40–60 Hz (one-way analysis of variance [1ANOVA], comparison between different frequencies, *F*_(11 108)_ = 10.91, *p* = 1.31 × 10^-12^, ***Figure 2a*** left). The overlapping proportions were the lowest in the same 40–60 Hz frequency range, compared with other frequencies (1ANOVA, *F*_(11 108)_ = 13.62, *p* = 1.01 × 10^-15^, ***Figure 2b*** left). We noted that at a low *C* value (0.16), the effects of stimulation frequency on both the persistence score (2ANOVA, *F*_(1 238)_ = 384.88, *p* = 8.82 × 10^-51^, ***Figure 2a*** right) and the overlapping proportion (2ANOVA, *F*_(1 238)_ = 220.49, *p* = 2.63 × 10^-35^, ***Figure 2b*** right) were much more prominent. Further detailed analysis showed that slow-gamma stimulations at 40–60 Hz were effective because they reduced the firing rates of non-cued memory neurons that were not being recalled by the cue input (***Appendix – figure 4***). For neural oscillations, we found that theta power was enhanced by rescue stimulations at a frequency of approximately 10 Hz, but was not significantly different from that by stimulation frequency in slow- and fast-gamma range (***Figure 2c***), but slow- or fast-gamma power was most effectively enhanced by stimulations at the same slow- or fast-gamma frequency (***Figure 2d-e***), as expected from previous experimental results (Etter et al., 2019). Overall, our results suggest that stimulations at the slow-gamma band are more effective for memory rescue than those at other frequencies.

**Figure 2.**
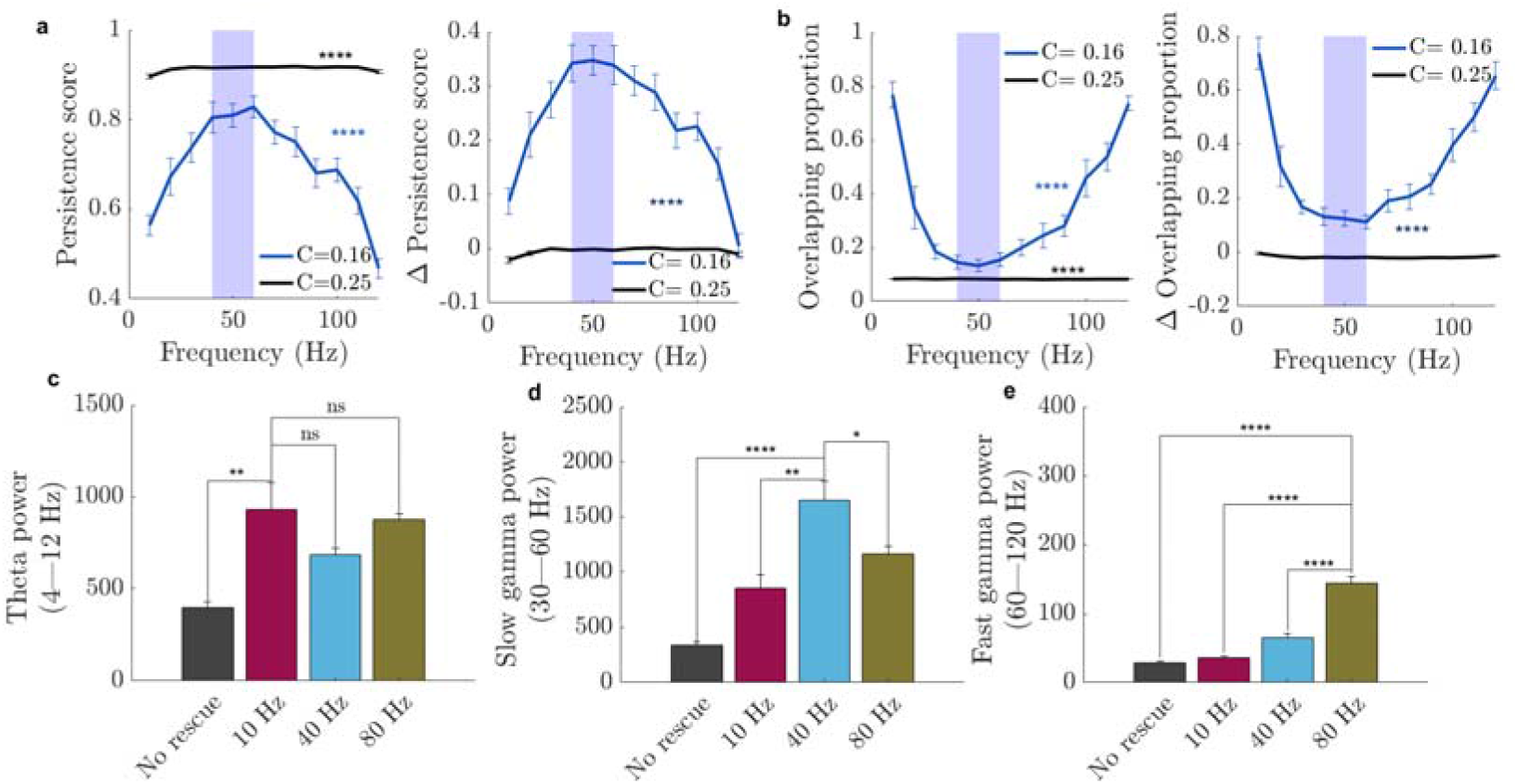
Rescue stimulations are more effective at slow-gamma frequencies. (a): Persistence score and (b): overlapping proportion and corresponding changes (effect size) after applying different rescue stimulation frequencies in the cases of high (0.25) and low (0.16) *C* values. The duty cycle was always set at 50%. The shaded area in (a, b) indicates the slow-gamma band. (c-e): Power of theta (c), slow-gamma (d), and fast-gamma (e) oscillations, with or without rescue stimulations, at different frequencies in the case of a low *C* value (0.16). Error bars: SEM. n = 10 trials/setting. In (a, b), one-way analysis of variance (1ANOVA) test; in (c-e), two-sample Student’s t-test. ns, *p* > 0.05; *, *p* < 0.05; **, *p* < 0.01; ****, *p* < 0.0001

### Learning a New Memory in the Presence of Existing Memories

In the work described in the previous sections, memory engrams were preset in the model to study memory recall and rescue. Here, we focused on how reduced *C* affects the learning of a new memory in the presence of preset memories. To this end, we considered a learning model (***Figure 3a***), consisting of nine preset memory engrams (neuron number 200–1,999) and one learning engram (neuron number 0–199) driven by a learning signal, which was a strong external input (12.5 Hz) applied for 100 s to the learning engram. The synaptic weights of neurons in the learning engram began at an initial non-coding value. When the learning signal was present, all excitatory synapses between all excitatory neurons (including those in the preset engrams) were dynamically changed according to a learning rule (see Methods) (Chistiakova et al., 2014; Kwon and Sabatini, 2011; Lev-Ram et al., 2002; Panda and Roy, 2017; Pfister, 2006; Zenke et al., 2015). Afterwards, learning was terminated (synaptic weights were fixed) and all preset and newly learnt memory engrams were sequentially recalled by applying the cue input one by one, as in our previous model (***Figure 1a***). We analysed how *C* affected learning-induced changes in the synaptic weights and recall activities of the newly learnt and preset engrams.

**Figure 3.**
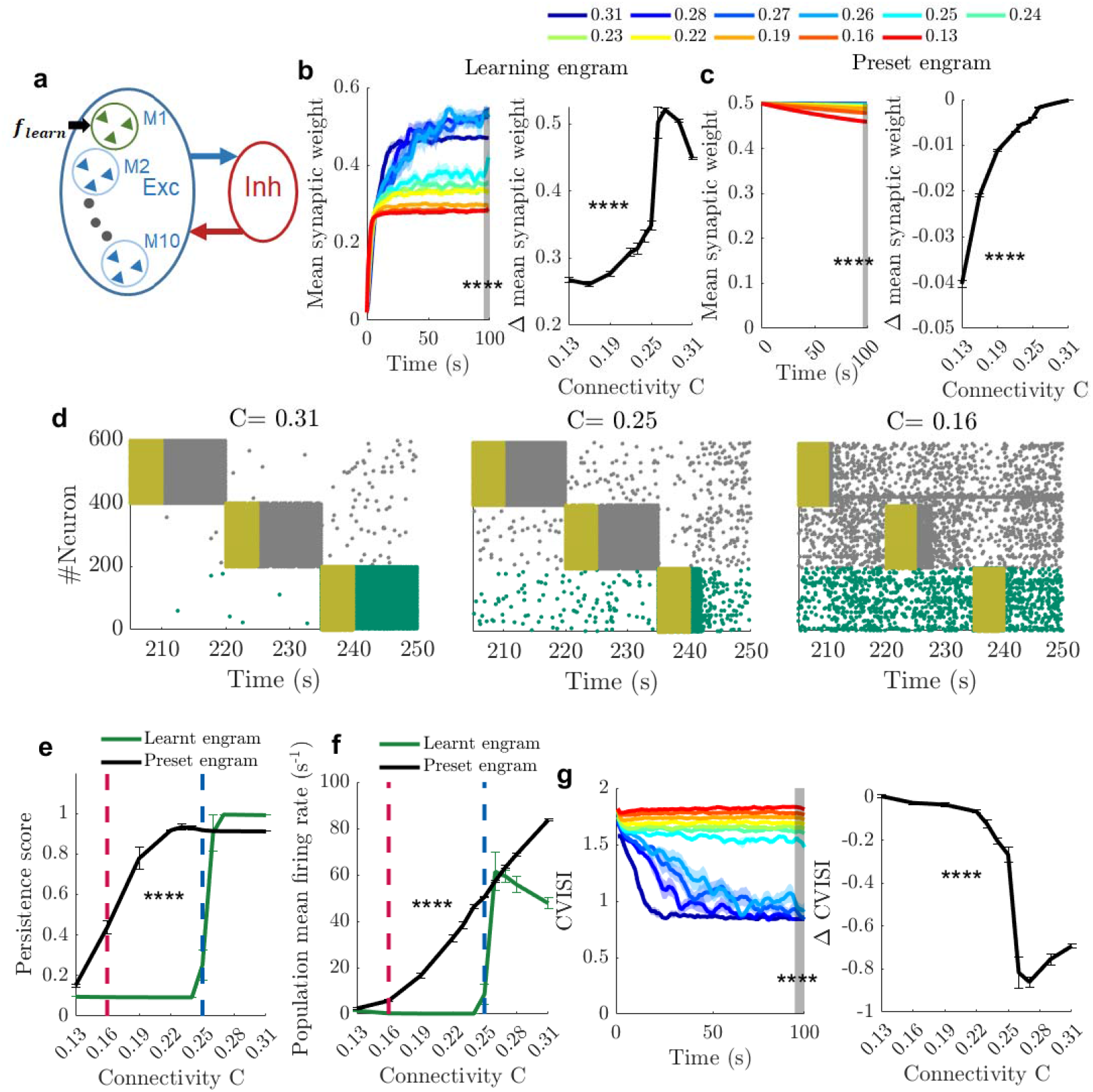
Low connectivity impairs the learning of new memory engrams. (a): Learning model with nine preset memory engrams (M2–M10) and a learning engram (M1) that is left for learning by receiving a learning signal *f_learn_*. (b-c): Evolution (left) and changes between 100 s and 1 s (right) of the mean synaptic weight within the learning engram (b) and preset engrams (c) during learning with different connectivities, *C.* (d): Spike raster plot of the learnt and two preset engrams during recall at different *C* values (C = 0.31, 0.25, 0.16). Grey/green: spikes of preset/learnt engrams, yellow: cue input. (e-f): Persistence score (e) and population mean firing rate (f) of the learnt engram (green) and preset engrams (black) at different C values during recall after learning. Vertical red and blue dashed lines indicate *C* = 0.16 and *C* = 0.25, respectively. (g): Evolution (left) and changes between 100 s and 1 s (right) of the coefficient of variation of the inter-spike interval (CVISI) within the learning engram during learning with different *C* values. Error bars in (E, F) and shading area in (b, c, g): SEM. ****, p < 0.0001 in 2ANOVA test; (e, f), the value at 100 s in 1ANOVA test (grey region in b, c, g).

We first examined how the synaptic weight evolves in the learning and preset engrams during learning depended on *C.* When *C* was large (> 0.25), the synapses between neurons targeted for learning tended to potentiate to a large value and form memory engrams, while when *C* was small (< 0.25) they tended to potentiate much less (1ANOVA comparing values at *t* = 100 s among different *C* values, *F*_(1 108)_ = 134.77, *p* = 5.11 × 10^-53^, ***Figure 3b*** left; 1ANOVA, _108)_ = 81.11, *p* = 5.69 × 10^-47^, ***Figure 3b*** right). In the preset engrams, the synapses were not affected by learning the new memory at large *C* values (> 0.25), and more synapses tended to depress as *C* was reduced, but the synaptic weight reduction was modest (1ANOVA test at t = 100s, *F*_(1 108)_ = 113.45, *p* = 4.76 × 10^-54^, ***Figure 3c*** left; 1ANOVA test, *F*_(1 108)_ = 2352, *p* = 0.00, ***Figure 3c*** right).

We then examined the memory recall of the learnt engram after learning at different *C* values. Examples of spike raster plots during recall showed that persistent states were successfully activated at a high *C* value (0.31) for the learnt engram (***Figure 3d***, left), whereas at small *C* values, they were either truncated or not activated (***Figure 3d***, middle and right). Our quantifications confirmed this observation. The persistence score and population mean firing rate of the learnt engram were high at large *C* values (> 0.25; ***Figure 3e, f***). Thus, in this regime, the new memory was successfully encoded. As *C* values were reduced (< 0.25), the persistence score and firing rate of the learnt engram decreased to almost zero, indicating an impairment in the encoding of the new memory. This encoding impairment was most likely caused by the low synaptic weights after learning at low *C* values (***Figure 3b***), because further analysis showed that synaptic strength within a memory engram needs to be sufficiently great for a cue input to produce persistent states during memory recall (***Appendix – figure 5***). In addition, we formally compared these measurements between the learnt and preset engrams during memory recall. Both measures were significantly lower for the learnt engram than the preset engrams, especially at low *C* values (persistence score: 2ANOVA, *F*_(1 238)_ = 58.81, *p* = 2.86 × 10^-28^, ***Figure 3e***; population mean firing rate: 2ANOVA, *F*_(1 238)_ = 39.36, *p* = 1.57 × 10^-9^, ***Figure 3f***). Our analysis thus indicates that low connectivity impairs the encoding of the learning engram, such that it cannot be properly recalled later as a preset engram.

We performed additional analysis to understand how the encoding impairment in the learning engram at low *C* values was produced during learning through the plasticity rule in our model. One clue came from our observation that at high *C* values, but not at low *C* values, neurons in the learning engram changed their firing burstiness during learning (***Figure 3g***), as quantified by the coefficient of variation of the inter-spike interval (CVISI), with a value of 1 indicating Poisson firing and a value > 1 indicating burst firing. At the beginning of learning, neurons in the learning engram were bursty (CVISI > 1.55) across all *C* values (driven by the learning signal). At the end of learning, however, they became much less bursty (CVISI close to 1) at high *C* values (> 0.25), but remained bursty at low *C* values (< 0.25) (1ANOVA, *F*_(1 118)_ = 103.84, *p* = 3.69 × 10^-52^; ***Figure 3g*** right gray region). Because synaptic weights within the learning engram also showed different dynamic changes at different *C* values during learning (***Figure 3b***), we probed a possible link between synaptic weights, firing rate, and CVISI in a simple two-neuron model, consisting of a presynaptic neuron and a postsynaptic neuron connected by a plastic synapse (see Methods). We found that after reaching a relatively high external firing rate during learning, strong burst firing (CVISI > 1.55) recruited heterosynaptic plasticity in the learning rule in our model, which restricted synaptic potentiation (***Appendix – figure 6***). Therefore, the sustained high-burst firing during learning at low *C* values presumably limited the synaptic potentiation within the learnt engram, leading to low synaptic weights and encoding impairment.

We finally examined the recall of preset engrams after learning. The preset engrams were generally recalled with higher persistence scores and population mean firing rates than the learnt engram (***Figure 3e, f***), as described above. In addition, the recall performances did not significantly differ from those without learning (***Appendix – figure 7***). Therefore, the modest weakening of synaptic weights within the preset engrams by learning (***Figure 3c***) did not further impair the recall of preset engrams, in addition to the impairment caused by low *C.*

Overall, our results showed that at a high level of connectivity (*C* > 0.25), a new memory engram can be learnt, whereas with reduced connectivity (*C* < 0.25), the encoding of the new memory engram was impaired. However, the recall of preset memory engrams was largely unaffected by learning.

## Discussion

In this work, we studied the effects of synaptic degeneration on the recall and learning processes in a neural network under different levels of connectivity. We first discovered that low *C* led to changes in the persistent state related to memory recall. Gradually reducing *C* reduced firing synchrony, but interestingly, increased then decreased the overall population mean firing rate of memory neurons. Second, we found that deactivating inhibitory neurons in the slow-gamma frequency effectively enhanced the firing rate and synchrony of cued memory engrams to partially rescue memory recall from engram activation impairment in low-*C* situations. However, low *C* may lead to the simultaneous activation of multiple memory engrams by slow-gamma rescue stimulations. Third, we studied the learning of a new memory engram in the presence of existing memory engrams. We found that with sufficiently high connectivity, the new memory engram can be successfully learnt. There was a mild reduction regime with impairment in encoding a new memory engram, but the activation of preset engrams was not affected by learning. There have been several simulation studies on the effect of synaptic loss. Abuhassan et al. studied the process of compensation to increase the excitability of neurons and reproduced the shift of the oscillation band to a lower frequency with synaptic loss (Abuhassan et al., 2014b, 2014a). In Ref. (Romani et al., 2013), the increased neurotransmitter release caused by Aβ (Abramov et al., 2009) was found to cause an increase in spiking probability, but a reduction in synaptic current generated by sequential spiking at different frequencies. In Ref. (de Haan et al., 2017), different therapeutic strategies were studied using a neural mass model with degenerated human network topology, and it was found that, although increasing the degree of inhibition by inhibitory neurons had favourable effects, the best strategy was to enhance the excitability of excitatory neurons. These studies have focused on the spontaneous dynamics of circuits with synaptic loss, but have not considered the existence and activation of engrams, which are important in learning and memory (Roy et al., 2016; Ryan et al., 2015); thus, our model has the advantage of filling this gap.

From a mechanistic point of view, memory deficits may be caused by a failure to form (encoding impairment) or activate (engram activation impairment) memory engrams. The results of previous experiments (Roy et al., 2016; Ryan et al., 2015) support an engram activation impairment underlying recall impairment in AD (Roy et al., 2016; Ryan et al., 2015; Stopford et al., 2007). In Ref.(Stopford et al., 2007), some AD patients showed poorer performance at recalling visual and verbal information immediately after a presentation and after a 30-minute delay. In this regard, our study shows that severe connectivity deficits induce recall difficulty in activating persistent states associated with previously encoded memory engrams. This is consistent with previous studies showing that persistent states are related to a bifurcation property (Chaudhuri and Fiete, 2016; Mongillo et al., 2008; Tsodyks, 2005) induced by changes in local synaptic weight or connectivity.

Experimental studies have shown that compared with apolipoprotein E (APOE) ε4 non-carriers, APOE *ε*4 carriers have higher brain activity in the early stage of life (Dennis et al., 2010; Filippini et al., 2011; Scarmeas et al., 2005), but the old pathological patient in the later stage of disease is characterised by lower brain activity (Borghesani et al., 2008; Filippini et al., 2011). Combined with previous findings (Li et al., 2009; Pascale et al., 2007; Shankar et al., 2007) that AD patients have a lower synaptic connectivity between neurons, we speculated that such a connectivity deficit (reduction in *C*) would first increase firing and then reduce firing as *C* further decreases. This speculation is supported by our model simulation results (***Figure 1f***) and is also consistent with the results of a recent study (Liang et al., 2021) showing that densely connected modular networks, compared with sparsely connected networks, have a lower population mean firing rate within modules through increased firing synchrony. The low population mean firing rate, but sufficient excitability, of circuits with sufficient density is helpful for supporting information processing ability in a normal, healthy primate brain, as a low firing rate is beneficial for metabolic energy saving and may reduce the risk of Aβ deposition, which is thought to be a by-product of metabolism waste (Dolev et al., 2013; Gerstner et al., 2012; Palop and Mucke, 2016).

There are reports that memory deficits may be rescued by several measures. Several experimental studies have shown that manipulations through chemical or optogenetic stimulation rescue spatial memory performance (Busche et al., 2015; Etter et al., 2019; Roy et al., 2016; Zheng et al., 2020). However, the mechanism underlying memory rescue is elusive. Based on the results of a previous experiment (Etter et al., 2019), in which spatial memory performance was rescued by applying optogenetic stimulations at a slow-gamma frequency to medial septum PV cells, we applied slow-gamma frequency stimulations to inhibitory neurons in our model and provided new insights into the mechanism of rescuing previously encoded memories. We revealed that the suppression of inhibitory neurons increased the firing of excitatory neurons in both cued and non-cued memory engrams. Thus, memory recall was partially rescued, but the overlapping proportion during memory recall was increased. However, there was an optimal stimulation frequency of approximately 40 Hz, as this frequency most effectively avoided the over-activation of non-cued excitatory neurons in low *C* situations, thus better preventing non-cued memory engrams from being activated spontaneously. Even so, we found that the rescue induced an increase in the overlapping activation of memory engrams as a side effect. This phenomenon has not previously been found experimentally and may be worth testing in future experimental studies.

Impairments in the formation of new memories during learning have been reported in AD (Cheng and Ji, 2013; Germano and Kinsella, 2005; Hodges et al., 1990; Stopford et al., 2007; Takashima, 2012; Weintraub et al., 2012). To study how synaptic degeneration affects learning impairment, our model used a composited plasticity rule, including spike-timing-dependent plasticity (STDP), heterosynaptic plasticity, and transmitter-induced plasticity, to study the engram formation ability under conditions of connectivity deficit. These plasticity forms identified in previous experiments (Chistiakova et al., 2014; Gerstner et al., 1996; Gilson et al., 2010; Kwon and Sabatini, 2011; Morrison et al., 2008; Panda and Roy, 2017; Song et al., 2000; Srinivasa and Cho, 2014; Zenke et al., 2015) are biologically plausible. The ability to form engrams using STDP has been known for a long time (Bayati et al., 2015; Srinivasa and Cho, 2014; Zenke et al., 2015). However, the traditional STDP rule suffers from explosive synaptic weight potentiation (Gerstner et al., 1996; Gilson et al., 2010; Morrison et al., 2008; Song et al., 2000). To prevent an explosion, different modification forms of STDP rules (Krunglevicius, 2016; Kumar and Mehta, 2011; Morrison et al., 2007) or additional plasticity rules (Maes et al., 2020; Zenke et al., 2015) were used. Among above, we used heterosynaptic plasticity, as previously described (Zenke et al., 2015). Previous experiments have shown that heterosynaptic plasticity is activated in response to a high neuronal firing rate (Chistiakova et al., 2014; Panda and Roy, 2017) to restrain synaptic potentiation. Therefore, this plasticity rule ensures that the synaptic weight does not explode at any neuronal firing rate. We also used transmitter-induced plasticity (Kwon and Sabatini, 2011; Zenke et al., 2015) in our study to prevent neurons from becoming silent during the simulation. Our model incorporating these biologically plausible plasticity rules suggested that mild connectivity deficits affect the formation of new engrams first, but do not strongly affect existing memory engrams. This may reflect the mechanism of new memory impairment found in AD patients (Cheng and Ji, 2013; Germano and Kinsella, 2005; Hodges et al., 1990; Stopford et al., 2007; Takashima, 2012; Weintraub et al., 2012). With a severe reduction of connectivity, the preset memory engrams cannot be activated after learning a new memory, which is consistent with the loss of long-term memory in AD patients (Cheng and Ji, 2013; Germano and Kinsella, 2005; Roy et al., 2016; Stopford et al., 2007).

The reduced ability to form new engrams is related to the increase in burst firing at low levels of connectivity, which induces stronger heterosynaptic plasticity to restrain the increase in synaptic weight while learning the new engram. This model prediction of the mechanism underlying impairments in learning new memories may need to be further examined in future experiments.

Furthermore, various types of plasticity and intrinsic property of neurons are affected by neural degeneration. Homeostatic plasticity has been found to be abnormal in AD patients (Jang and Chung, 2016; Megill et al., 2015). Moreover, Aβ concentration has been found to affect the activation of *N*-methyl-d-aspartate (NMDA) receptors (Palop and Mucke, 2010; Puzzo et al., 2008), which is crucial for STDP (Bi and Poo, 1998; Kim et al., 2016; Poo et al., 2016). The activation of NMDA receptors in AD patients is potentiated in the early stage of disease and reduced in the late stage of disease (Palop and Mucke, 2010). Abnormal NMDA receptor activation in AD would, thus, lead to impaired plasticity. Lastly, the excitability of neurons reduces as aging and AD(Maurer et al., 2017; Tamagnini et al., 2015). AD plaque was found reduce inhibition of nearby synpases(Busche et al., 2008). The effect of homeostatic plasticity, impaired plasticity itself and the excitability changes warrant inclusion in further studies of connectivity deficits

Overall, our modelling work sheds light on the changes in memory engram dynamics caused by connectivity deficits related to neurodegeneration. The specific changes revealed here may inspire new experiments to examine different aspects of memory deficits in memory recall or learning.

## Methods and methods

### Network and Circuit Model

The model circuit we used was composed of 2,400 neurons, with 2,000 excitatory (E), *n_E_,* and 400 inhibitory (I), *n_I_*, neurons (***Figure 1a***). Neurons were connected randomly with directed synapses with various probabilities, *C.* Four hundred independent Poisson trains with a rate of *f_background_* = 2.5 Hz were applied to each neuron to mimic the input from other neural circuits. The neural dynamics in the network were modelled by integrate-and-fire neurons with α-amino-3-hydroxy-5-methyl-4-isoxazolepropionate (AMPA)-, NMDA-, and gamma-aminobutyric acid (GABA)-receptor-mediated conductance (Yang et al., 2017) as follows:

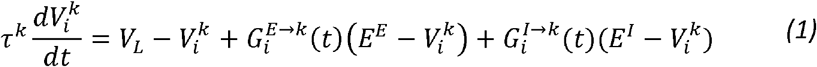

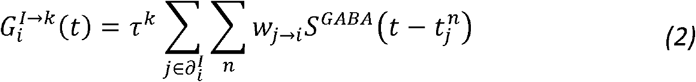

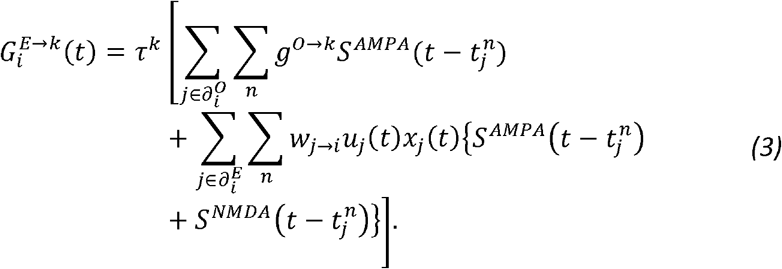

Here, *k* = *E* or *I* denotes the neuron type and 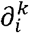 indicates the neighbours with *k* neuron type of neuron *i. O* → *k* indicates the external input to neuron 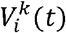 is the membrane potential of neuron *i* and neuron type *k*, and 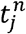 is the *n*-th spike of neuron *j*. The synaptic time course is described by the following bi-exponential function:

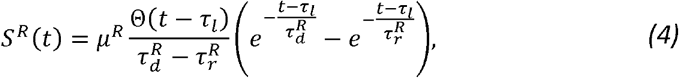

where 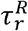 and 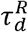 are the characteristic rising time constant and decay time constant, respectively, and *μ^R^* is the amplitude, depending on the receptor type *R* (AMPA, GABA, or NMDA) of the presynaptic neurons. Θ(*t*) is the Heaviside step function. *w_j→i_* is the synaptic weight from the presynaptic neuron, *j*, to the postsynaptic neuron, *i*, and was initially set as

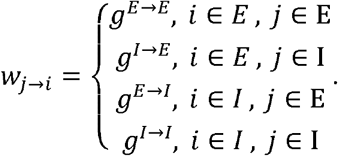

Two hundred excitatory neurons were randomly assigned to an engram representing a memory and there were several non-overlapping memory engrams considered in our model. The synaptic weights within each engram were replaced from the baseline value, *g^E→E^*, to a larger value, 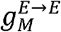.

### Short-Term Plasticity

Presynaptic short-term plasticity was described by the dynamics of two variables, the amount of neurotransmitter, *x_i_*, and the release probability, *u_i_*, in Eq. (3), and was governed by a depression time constant, *τ_D_*, and a facilitation time constant, *τ_F_,* as described in the following equations (*τ_F_* » *τ_D_*):

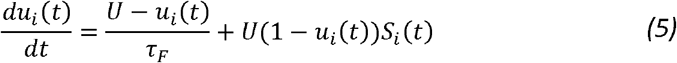

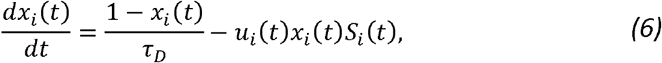

where 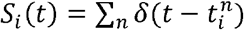 is the spike train of neuron *i* and *δ*(*t*) is the Dirac delta function. Depending on the firing history of a neuron, short-term depression or potentiation (Mongillo et al., 2008; Wu et al., 2020; Zucker, 2002) may be induced by the depletion of neurotransmitter, *x_i_*, or an increase in the release probability, *u_i_*, which temporarily changes the synaptic conductance, 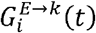, in Eq. (3).

### Long-Term Plasticity

The long-term change in synaptic weight obeys an integrative plasticity rule, which includes triplet STDP, heterosynaptic plasticity, and transmitter-induced plasticity (Zenke et al., 2015). The plasticity rule was applied to all E-to-E synapses. The synaptic weight evolved according to the following formula:

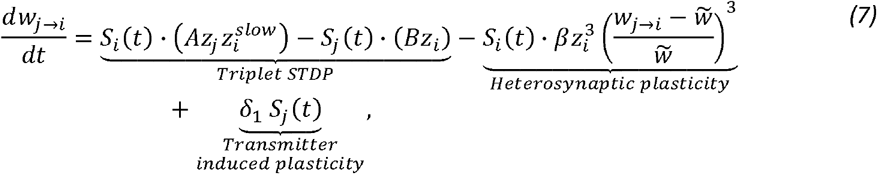

where *j* → *i* indicates the connection from neuron *j* to neuron *i.* The synaptic traces evolved as follows:

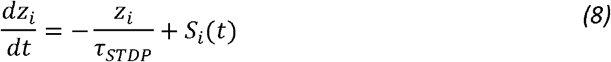

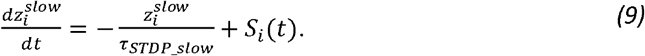

Synaptic weight modification was controlled by the presynaptic trace, z_j_ and the postsynaptic traces, *z_i_* and 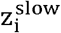, which increased by one with every spike and decayed to zero with the characteristic time constants, *τ_STDP_* and *τ_STDP_slow_,* respectively. The slow synaptic trace, 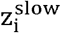, kept the spiking information of the long timescale *τ_STDP_slow_,* which made the synaptic potentiation term of the triplet STDP, 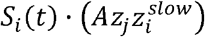, depend on the postsynaptic spiking rate and increased the synaptic weight by the amount 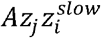 with every spike. The depression term of the triplet STDP, –*S_j_*(*t*) · (*Bz_i_*), reduced the synaptic weight by the amount *Bz_i_* with every presynaptic spike. The triplet STDP rule alone was unstable because potentiated (depressed) synapses tend to induce more (less) postsynaptic spikes, which in turn potentiate (depress) the synapses. To stabilise synaptic weight, heterosynaptic plasticity, the third term in Eq. (7), was introduced in (Zenke et al., 2015). Heterosynaptic plasticity reduces the synaptic weight by the amount 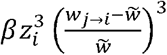, where 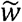 is the preferred synaptic weight at every postsynaptic spike to prevent synaptic weight explosion. Heterosynaptic plasticity has been identified in previous experiments (Chistiakova et al., 2014; Panda and Roy, 2017) and is believed to occur in the high-firing-rate domain. Therefore, we modelled heterosynaptic plasticity as the third power of the postsynaptic trace, 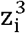. Transmitter-induced plasticity, the last term in Eq. (7), prevents neurons from being silent, reflecting spontaneous spine growth (Kwon and Sabatini, 2011) or a potentiation mechanism that counteracts Hebbian long-term depression (Lev-Ram et al., 2002). In addition, according to Dale’s principle, a lower bound of 0.001 was set for each plastic synapse to prevent negative and zero weights. The parameters of the model and the plasticity rules are shown in ***Appendix — Table 1***.

### Simulation of Memory Recall, Rescue, and Learning

From the beginning of the simulation, each neuron received 400 independent Poisson pulse trains at 2.5 Hz as the background external input to mimic the inputs from other neural circuits, which were present during the entire simulation time. The simulation was first run for 5 s to avoid a transient state. When studying memory recall, memory engrams were activated in sequence by adding an additional external cue input of 10 Hz, *f_cue_*, to one engram for 5 s. The next memory engrams were then cued consecutively after 10 s (***Figure 1b***). Therefore, the total time interval between the start of the cue of consecutive memory engrams was 15 s.

To study memory rescue, rescue stimulations, with a duty cycle of 10%–90%, were applied to 50% of the inhibitory neurons randomly selected at the beginning of the simulation, at a frequency of 10–120 Hz. A duty cycle of 10%–90% corresponds to on-stimulus for the first 10%–90% of time in each cycle of the stimulation and off-stimulus for the remaining time. When the stimulus in the rescue stimulations was on, the membrane potential of each chosen inhibitory neuron, if it was not within the refractory period, had a 50% chance per time step to be reset to its leakage potential, V_L_, which prevented the chosen neurons from firing at the moment of applying the rescue stimulation.

When studying the learning of a new memory in addition to existing memories at different *C* values, nine engrams were preset and one additional engram (neuron number 0–199) was left for learning (learnt engram). The synaptic weights between neurons in the learning engram started at an initial non-coding value of *g^E→E^* = 0.02. After a 5 s period of simulation with the background input, the learning process began and lasted for 100 s, during which Poisson learning inputs at 12.5 Hz were applied to the learnt engram in addition to the background input. In the learning period, all synaptic weights between the excitatory neurons, including those in the preset engrams, were updated according to the learning rule described above (Eq. [7]). We tuned the learning rule parameters such that the distribution peak of the synaptic weight after learning was located at approximately 0.5 to 0.6 for high *C* values (***Figure 3b***), which is comparable to the average, 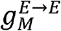, within the preset engrams. After learning, changes in synaptic weight were prohibited and all preset and new memory engrams were cued one by one for memory recall.

### Simple Model of Two Connected Neurons

To gain insights into how synaptic weights change with different presynaptic and postsynaptic firing rate and firing pattern combinations, we built a simple two-neuron model with one synapse to learn using the learning rule described in Eq. (7). The parameters used were the same as those listed in Supplementary Table 1. The simulation lasted for 100 s and the firing activities of the two neurons were generated from a Poisson random process with fixed rates that did not change with learning. Therefore, in this model, the causality of presynaptic firing increasing the firing of postsynaptic neurons and the effect of synaptic weight change during learning on firing dynamics were ignored.

In addition to firing rate, the bursty firing of neurons, which was represented by the CVISI (higher CVISI, more bursty), also affected the synaptic weight. To study the effect of the CVISI on synaptic weight change (under a fixed total number of spikes), the spike trains of the presynaptic and postsynaptic neurons used in the two-neuron model were manipulated to introduce burst firing. *N* spikes were randomly chosen with a probability *p_pick_*, and their corresponding spike times were 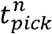, *n* = 1,2, …*N*. Two new spikes were added after every picked spike, at 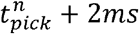 and 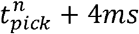, to form a new spike train. 2*N* spikes were then removed randomly from the new spike train to maintain the same number of spikes (*N* spikes; see illustration in ***Appendix – figure 6C***). Under this manipulation, the CVISI increased when *p_pick_* increased, as more spikes were reorganised to turn into the burst firing of 2–3 spikes.

### Simulation Details and Code

All simulation programs were written in C++ and run on a computer cluster supported by the High-Performance Cluster Computer Centre at Hong Kong Baptist University. The equations for the circuit model and plasticity rules were simulated using the second-order Runge–Kutta method, with a time step of 0.05 ms and the spike time correction method, as previously reported (Shelley and Tao, 2001).

### Firing Properties

We quantified the firing properties of neurons in cued-memory engrams (memory neurons) and outside the engrams (non-cued memory neurons). When studying the persistent state, we computed the population mean firing rate, which is the number of spikes fired by all neurons in an engram in a time window of 10 s from the end of their cue input, divided by the window time and the number of neurons in the engram. Moreover, when studying the learning process, we computed the firing rate distribution for the last 20 s of learning.

In addition, we computed the proportion of high-firing neurons in the engram, defined as those with an individual firing rate larger than 5 *s*^−1^, which is the firing threshold separating the low-activity state from the persistent state, as described in **Memory Recall** in **result,** from the cue input termination to 10 s after termination.

To quantify the synchronisation of neurons, the synchrony index, *K_ij_*, which is a measure of the probability of a pair of neurons firing coincidently, was defined as,

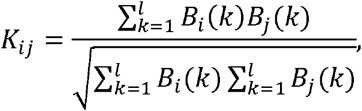

where *B_i_*(*k*) and *B_j_*(*k*) indicate neuron *i* and *j*, respectively, with a spike at the *k*-th 1 ms bin (0 for no spike and 1 for having a spike). A high synchrony index indicates that the neurons tend to fire coincidently.

### Persistent States, Persistence score, and Overlapping Proportion

To study memory recall, we detected cue-induced persistent states for each memory engram. A persistent state of an engram is defined as the period with a population mean firing rate larger than 5 s^-1^, to separate the persistent state from a low-activity state, which is also valid when implementing rescue stimulations, detected using a moving window of 1 s with a time step of 1 ms. If a gap with a low population mean firing rate (< 5 s^-1^) lasting for less than 1 s appeared between two persistent states, the gap was ignored and the two persistent states were counted as one persistent state.

The firing rate threshold was determined from the firing rate distribution of all excitatory neurons under different conditions (***Appendix – figure 2***). It was calculated using a 1 s moving window with a 1 ms time step. The results for different moving windows from different neurons were pooled when calculating the distribution. There were two far-separated groups corresponding to low-activity and persistent states, which had firing rates less than 5 s^-1^ and much larger than 5 s^-1^, respectively. Therefore, we chose 5 s^-1^ as the firing rate threshold, but as the separation was wide, there was a broad range of options for choosing the threshold at which the separation result shows little change.

We computed a persistence score for each persistent state, assuming that a persistent state activated by the cue input to one memory engram ideally persists until it is stopped by the cue input to another memory engram, as described in the following formulae:

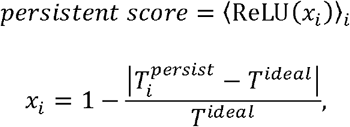

Where 〈·〉_*i*_ is the mean of neurons in the memory engram, *i*; ReLU is the rectified linear function, expressed as ReLU(*x*) = max(0, *x*); 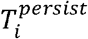 is the total duration of the persistent state of the memory engram, *i*, excluding the cue input period; and *T^ideal^* is the time difference between the termination of the cue input for one memory and the onset of the cue input for another memory. If the persistent state lasted for *T^ideal^*, the persistence score was 1. If the persistent state was different from *T^ideal^,* either shorter or longer, the persistence score was less than 1. A smaller score indicated that 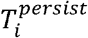 was further away from *T^ideal^.*

To quantify the degree of co-activation of multiple memories, we computed the overlap among the persistent states of all memory engrams using an overlapping proportion, defined

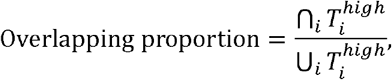

as where 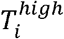 is the total time of the persistent state for the memory engram, *i*, excluding the cue input period. The overlapping proportion was bounded by 1, and a larger value indicated more co-activation of multiple memories.

### Oscillation Power

To quantify the oscillation in the simulation, we calculated the power spectrum using the fast Fourier transform of the mean-detrended average membrane potential of all excitatory neurons in the 10 s time interval after the cue input termination and normalised it to the mean of the power spectrum.

## Appendix

**Appendix – figure 1.**
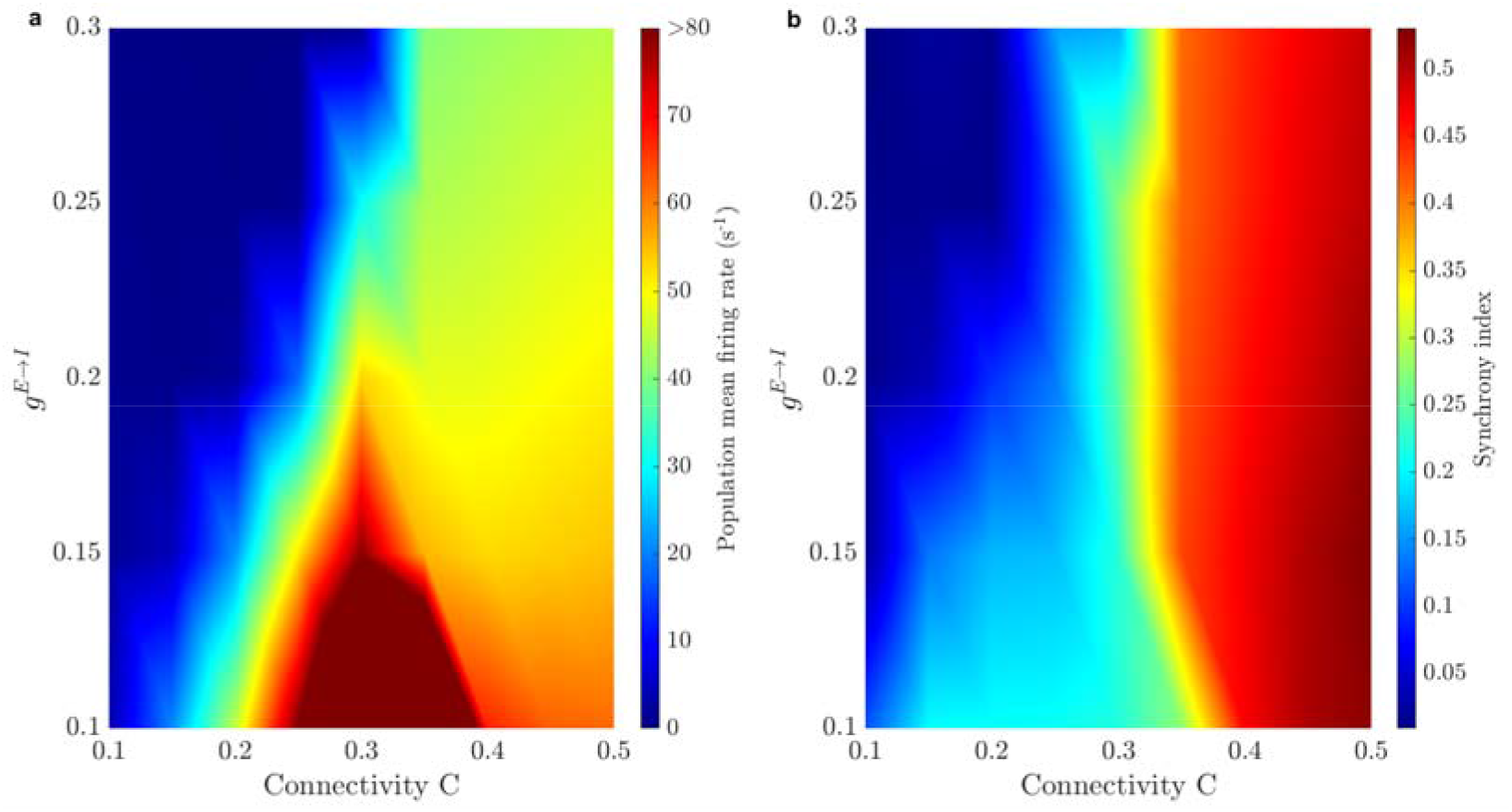
Cue-induced firing properties of memory neurons change with connectivity *C* in a wide range of values for the synaptic weight between excitatory and inhibitory neurons *g^E→I^,* in the circuit in Figure 1 without rescue stimulations. (a): Population mean firing rate. (b): Synchrony index calculated in a time window from the cue termination to 10s afterwards. The increase in population mean firing rate when *C* reduces happens when *g^E→I^* ranges from 0.1 to 0.2 and when *C* < 0.2, firing rates and firing synchrony are low across all *g^E→I^,* suggesting a robust loss of persistent states.

**Appendix – figure 2.**
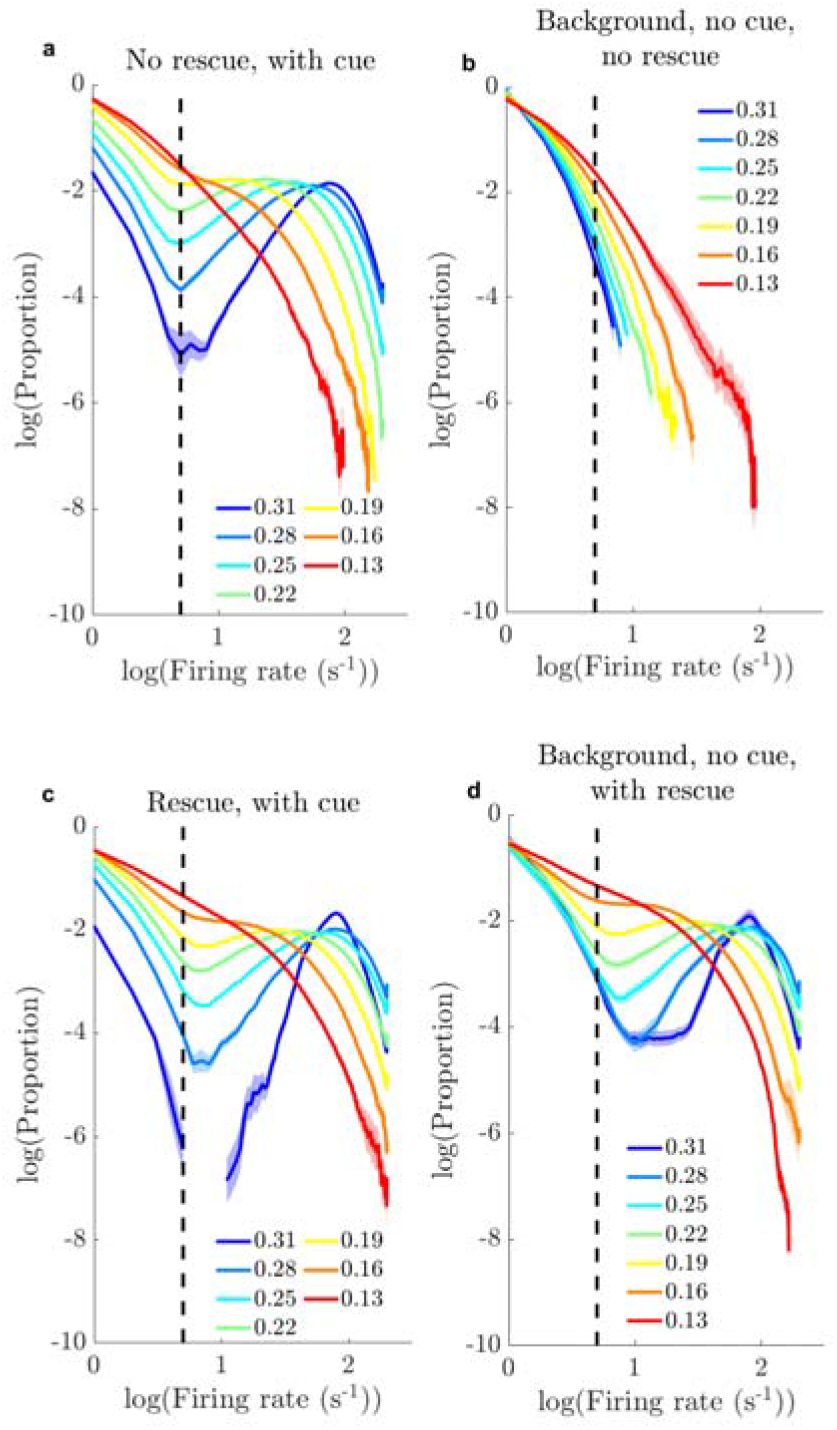
Distribution of the firing rate of excitatory neurons in the network under different conditions. (a): Sequential recall of memory engrams (within 10s after the termination of the cue input). (b): Background activity in the absence of cue input. (c): Same as (a), but with rescue stimulations (40Hz 50% duty cycle). (d): Same as (b), but with rescue stimulations (40Hz 50% duty cycle). The zero proportion in (c) at *C* = 0.31 causes undefined value and thus the curve is separated into two. Vertical dash line indicates firing rate at 5 *s*^-1^. n=10 trails for each *C* value. Shading area: SEM. Note the log scale on x and y-axis. In (a, c), the distribution of firing rate of excitatory neurons displays a bi-modal distribution, suggesting a distinction between a background state with low rates and a persistent state with high rates, separated at ~5 *s*^-1^. In (d), one memory engram may be activated spontaneously when *C* is high in the absence of cue and rescue stimulations.

**Appendix – figure 3.**
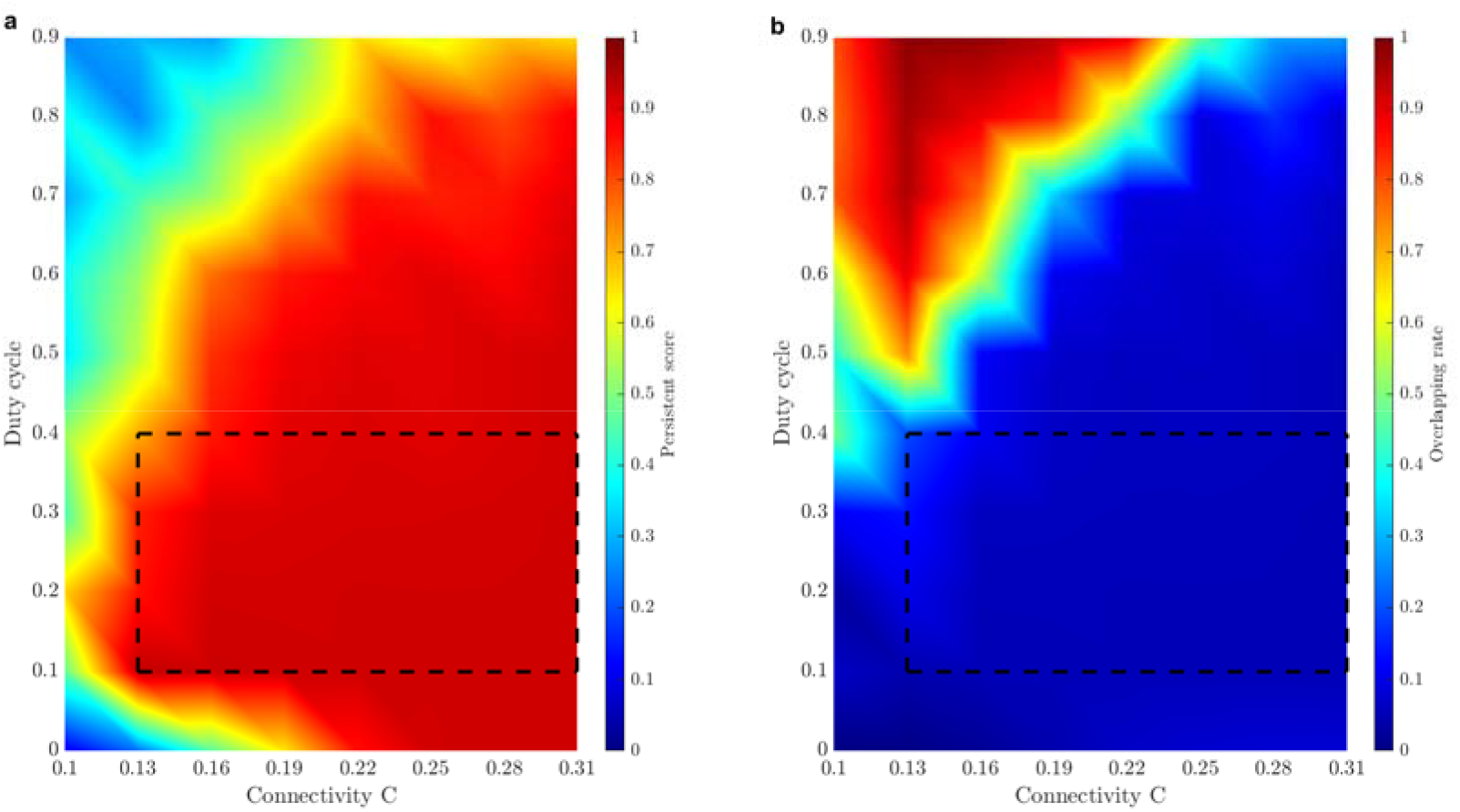
Quantifications of persistent state at different connectivity *C* with different duty cycles of slow-gamma stimulations (at 40Hz). (a): Persistence score. (b): Overlapping proportion. Dash rectangles indicate the duty cycle range with high persistence score and low overlapping proportion when *C* > 0.13. The result suggests that memory rescue by slow-gamma oscillations may be improved when the duty cycle of optogenetic stimulation is reduced (50% is used in the main text).

**Appendix – figure 4.**
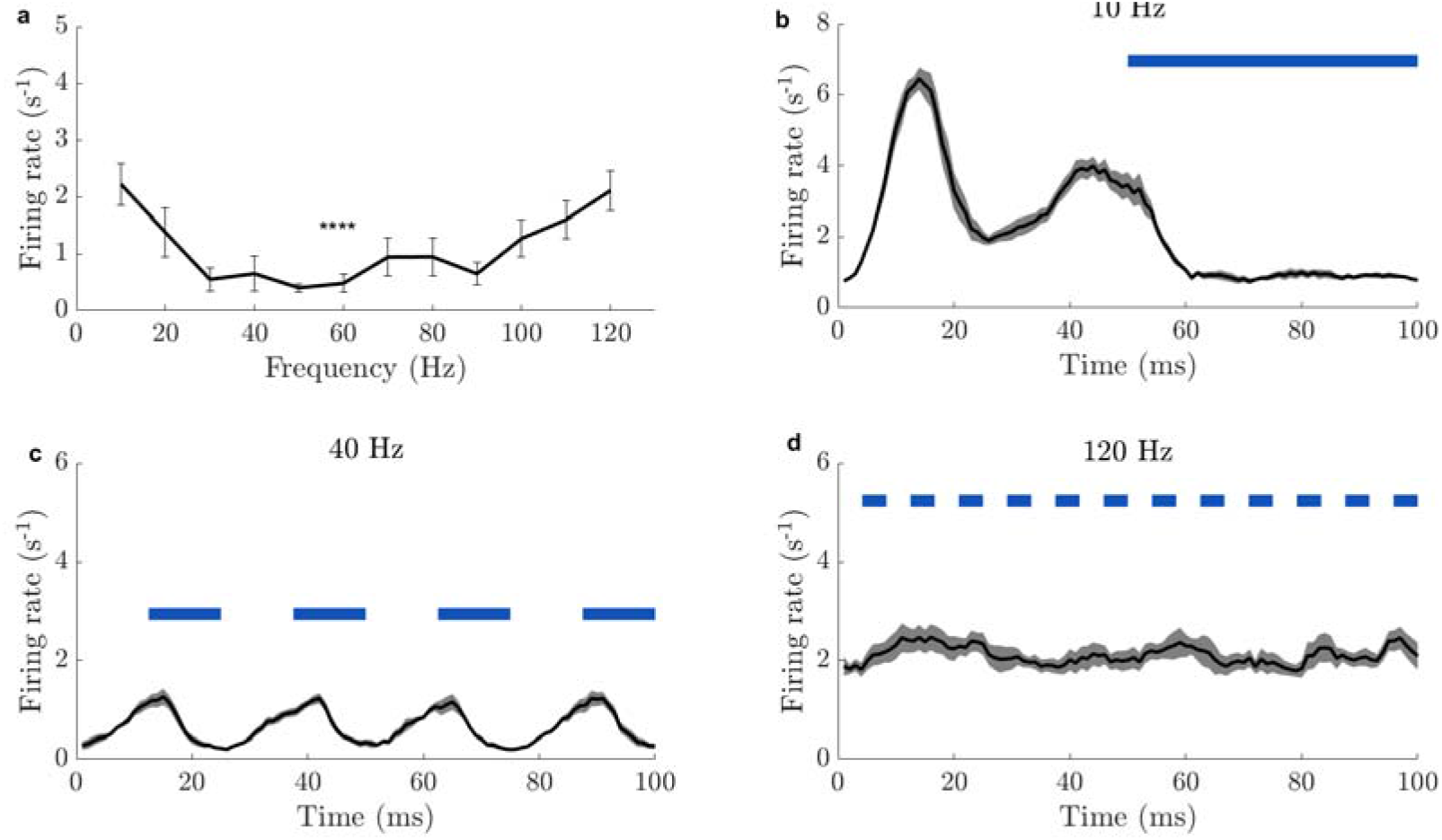
Firing rate of non-cued memory neurons at different rescue stimulation frequencies. (a): Population mean firing rate of non-cued neurons at different frequencies. It shows the population firing rate is minimal in the slow-gamma range. (b-d): Average population mean firing rate of non-cued neurons during the periods when rescue stimulations are applied. Broken blue bars indicate the on-off duty cycle with different frequencies: 10Hz (b), 40Hz (c), 120Hz (d). When the stimulation frequency is too low (10Hz), the population mean firing rate in the off-period increases. If the frequency is too high (120Hz), the population mean firing rate in the off-period does not respond to rescue stimulations and stays at high value. Only when the frequency is in a suitable range (around 40Hz), the population mean firing rate in the off-period can be suppressed effectively. Error bars in (a) and shading area in (b-d): SEM. N=10trials/setting. ns, *p* > 0.05; *, *p* < 0.05 in 1ANOVA test.

**Appendix – figure 5.**
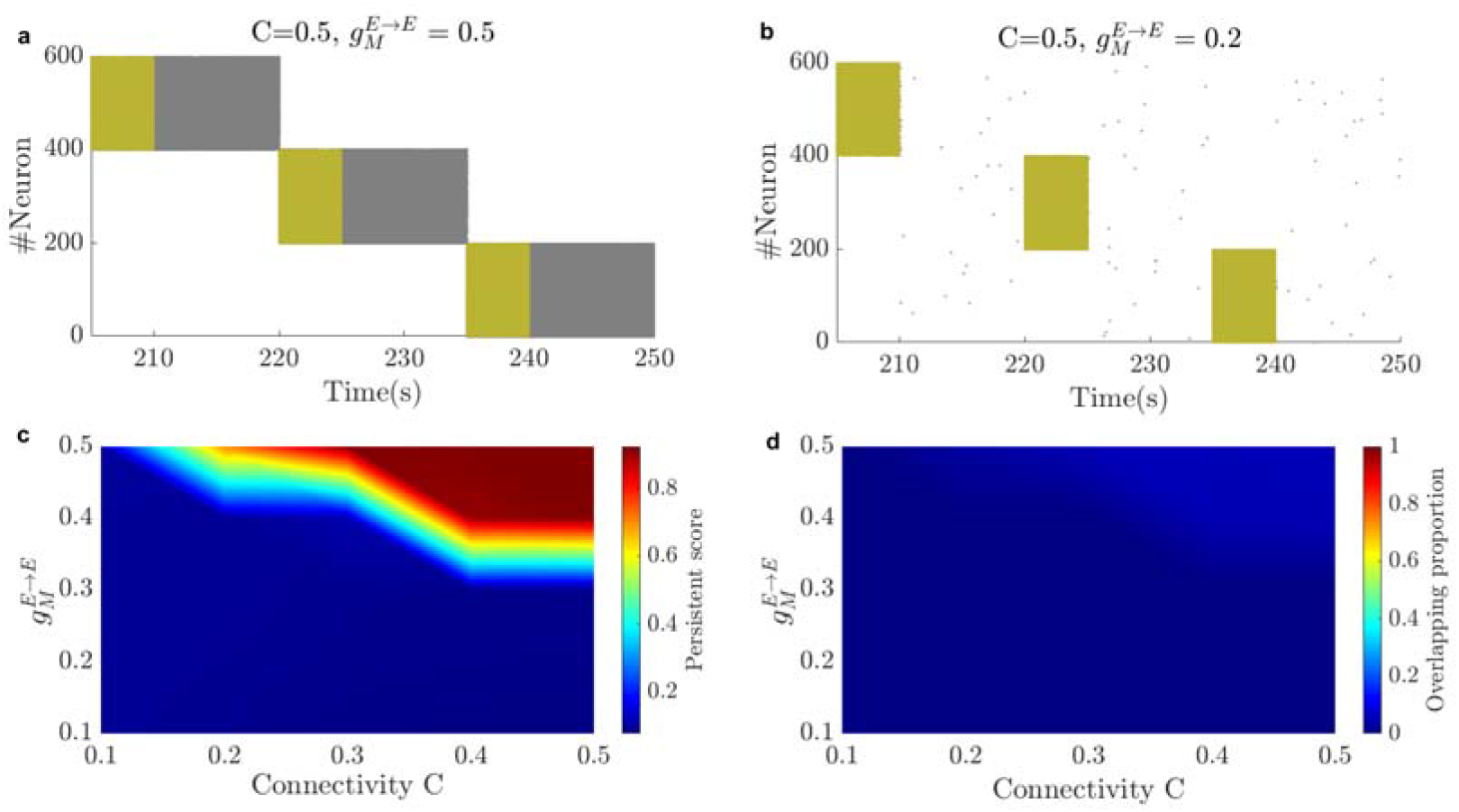
Persistent state at different connectivity *C* and at different within-engram synaptic weights 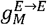. (a, b): Raster plot of *C* = 0.5 in the circuit in Figure 1A (without rescue) at (a): 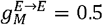; (b): 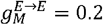. The periods with the strong cue (12.5Hz) input (5s) are marked in yellow. (c): High *C* and 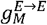 induces high persistence score. (d): The overlapping proportion is low among settings of different *C* and 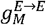. The persistence score at a given *C* drops to a value close to 0 as 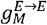 is reduced. Therefore, persistent states can only be activated when synaptic weights within engrams are sufficiently large.

**Appendix – figure 6.**
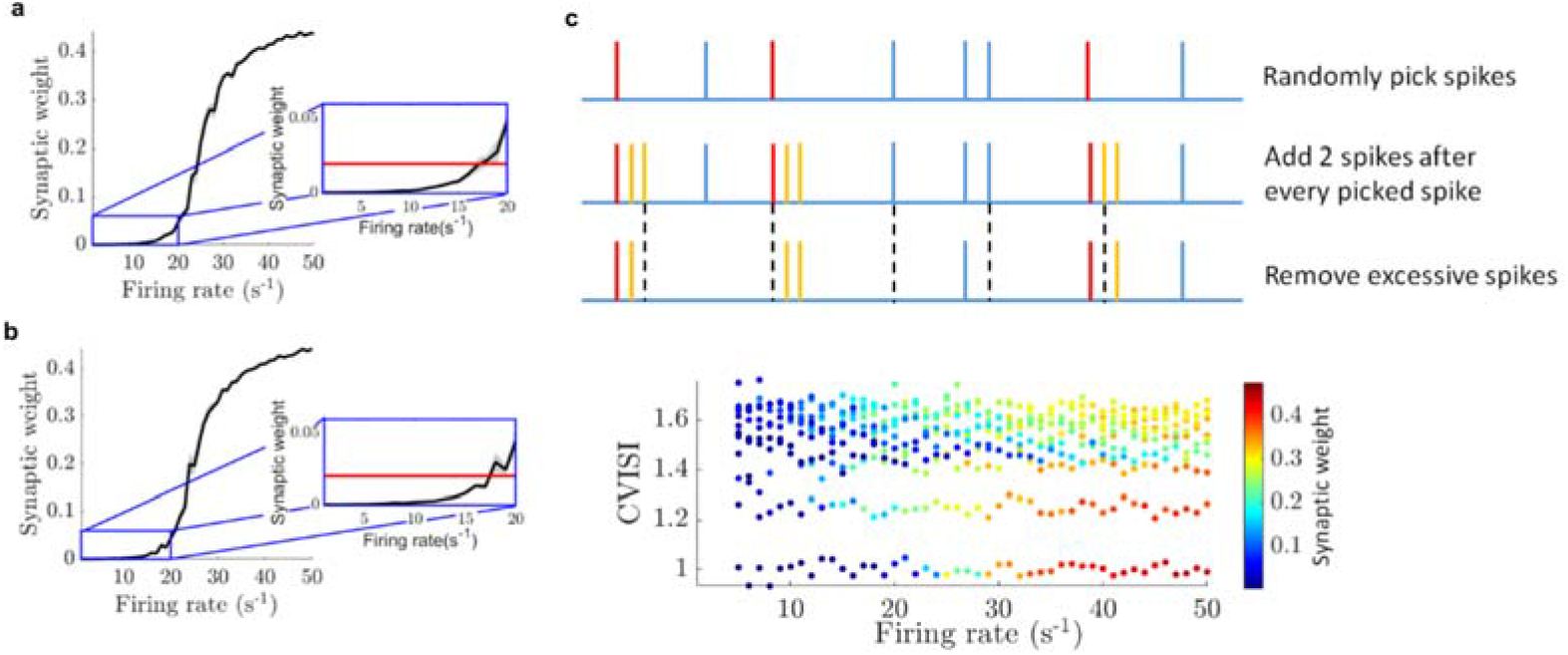
Dependence of final synaptic weight after learning on firing rate and CVISIs in a 2-neuron simple model. (a): Dependence of the synaptic weight (black line) on the firing rate (the same for presynaptic and postsynaptic neurons), with CVISI=1 (Poisson spike train). Insert plot is a magnification plot of postsynaptic neuronal firing rate arranging from 0 to 10*s*^-1^. Red line in the insert indicates the synaptic weight before learning. It shows when post-synaptic firing rate, is larger (smaller) than around 10*s*^-1^, the synapse is potentiated (depressed). n=10 trials/setting. Shading area: SEM. (b): Dependence of the synaptic weight (black line) on the postsynaptic neuronal firing rate, with the presynaptic neuronal firing rate fixed at 50*s*^-1^ and CVISI=1. Insert plot is a magnification plot of postsynaptic neuronal firing rate arranging from 0 to 10*s*^-1^. Red line indicates the synaptic weight before learning. n=10 trials/setting. Shading area: SEM. (c): (Top): Illustration of spike train generation to manipulate CVSI. 1) Pick spikes randomly (red lines) with probability *P_pikc_* from Poisson spike train. 2) Add 2 spikes (orange lines) after every picked spike. 3) Remove spikes randomly (black dash lines) to maintain the number of spikes as the original Poisson spike train. (Bottom): Synaptic weight (color coded) depending on the firing rate (the same for presynaptic and postsynaptic neurons) and CVSI. Change of CVISI is obtained by spike train manipulation in (b). The difference in synaptic weights is because large CVISI implies there is temporally high firing rate, which would activate the heterosynaptic depression to restrict potentiation by triplet synaptic plasticity. The dependence on CVISI is more important than the dependence on firing rate in the study of learning because the firing rate is at high value (>30 Hz) during learning.

**Appendix – figure 7.**
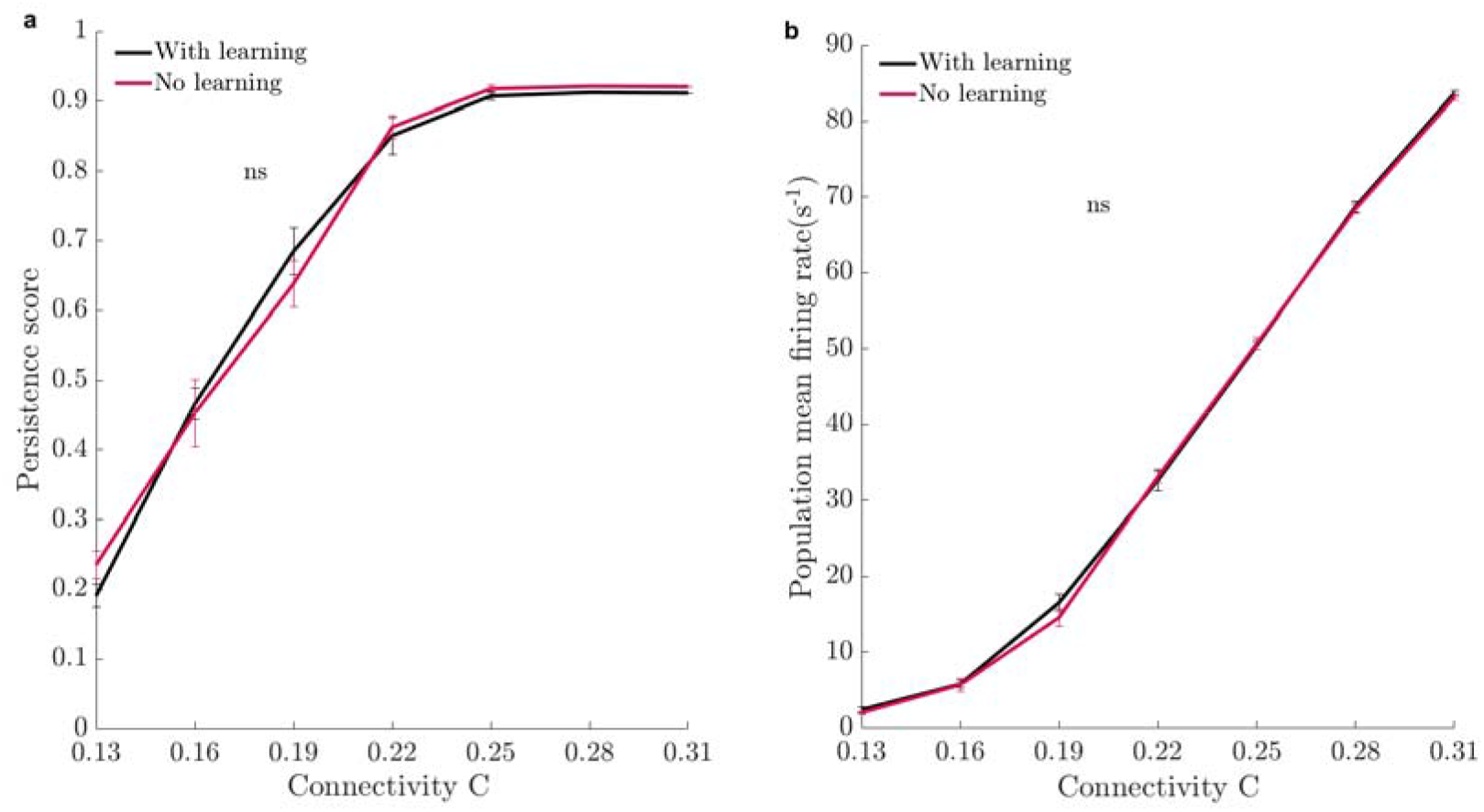
Preset engram deterioration. (a): Persistence score and (b): Population mean firing rate of preset engrams during recall with (Black) and without (red) learning before recall. n=10 trials/setting. Error bars: SEM. ns, *p* > 0.05 in 2ANOVA in (a, b). Although synaptic weights in the preset engams are modestly impacted by learning (Figure 3C), the effect is not strong enough to further impair the recall of preset engrams

**Appendix — table 1.**
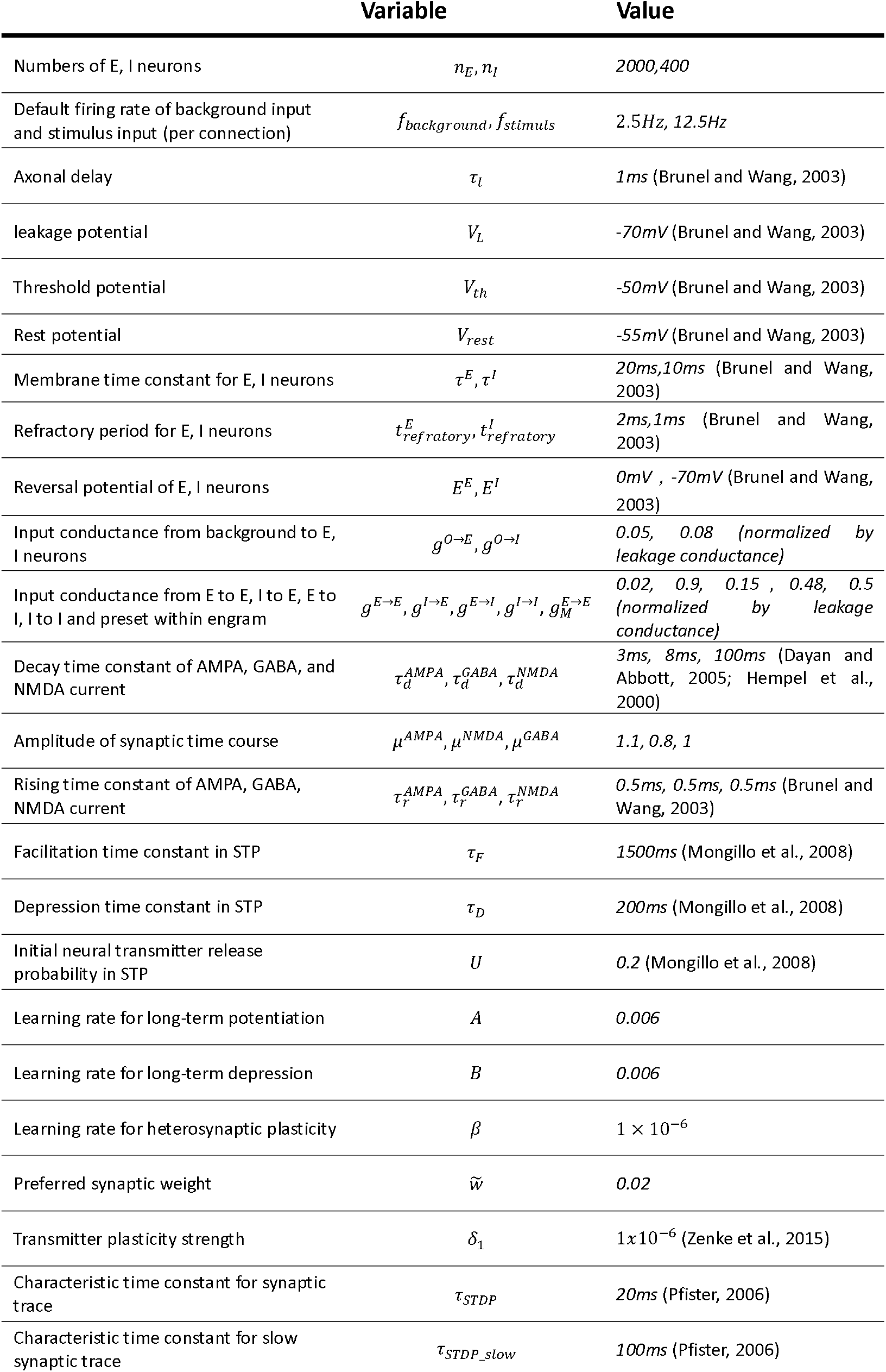
Parameters used in the neural circuit.

## Notes

### Competing Interest Statement

The authors have declared no competing interest.

